# Integrated deep learning and geo-referencing for drone-based animal tracking with flexible camera angles

**DOI:** 10.1101/2025.02.05.636599

**Authors:** Imran Samad, Dipani Sutaria, Damien R. Farine, Kartik Shanker, Mauricio Cantor

**Affiliations:** Centre for Ecological Sciences, Indian Institute of Science, Bangalore; Dakshin Foundation, Bangalore; School of Marine Biology and Aquaculture, James Cook University, Australia; Division of Ecology and Evolution, Research School of Biology, Australian National University, 46 Sullivans Creek Road, Canberra, ACT 2600, Australia; Department of Evolutionary Biology and Environmental Science, University of Zurich, Zurich, Switzerland; Department of Collective Behaviour, Max Planck Institute of Animal Behavior, Konstanz, Germany; Department of Fisheries, Wildlife and Conservation Sciences, Marine Mammal Institute, Oregon State University; Newport, OR 97365, USA

**Keywords:** Unmanned aerial vehicles, deep learning, geo-referencing, animal tracking, group tracking, behaviour

## Abstract

The ability of drones to provide detailed information on animals and their surroundings makes them ideal for studying animal behaviour at fine scales. While drones can provide high-resolution images of what animals are doing, they should also, in theory, be able to provide data on where they are. However, reconstructing geo-referenced tracks from drone videos that follow animals is challenging, particularly because current methods require specific drone flight patterns and large computational power.
Here, we combine deep learning and object tracking methods with a novel geo-referencing algorithm which allows us to track individuals across video frames and reconstruct their geo-referenced trajectories. We used a Region-based Convoluted Neural Network to detect animals and a Hungarian tracking algorithm to link detections across video frames, and then geo-referenced each detection in every frame to reconstruct individual trajectories.
We tested our geo-refencing algorithm through multiple drone flights with varying flight parameters over known Ground Control Points. The median (95% CI) geo-referencing error was 2.81 (0.74 – 23.23) meters, which reduced by 50% when the drone camera was positioned between -90° and -40°. Error increased with drone height and camera angle (-90° refers to the camera pointing towards the ground) but was not impacted by drone orientation.
We then demonstrate the utility of our framework with empirical examples using consumer-level drones. First, we tracked a volunteer carrying a high-resolution GPS unit and overlayed their GPS tracks on our estimated tracks to quantify tracking error. Next, we used drone videos of two delphinid species (*Tursiops truncates gephyreus*, and *Sousa plumbea*) representing varying environmental and flight conditions. We were able to successfully infer individual tracks across all conditions except when individuals formed tight clusters, in which case tracks were assigned a group identifier.
Our framework demonstrates an easy and robust approach to translate drone videos of moving animals into geo-referenced animal tracks which is applicable in many research contexts. A major advance over previous methods is that our algorithm is robust to different camera angles, and provides tracks with accuracy on-par with, or even exceeding, the accuracy from GPS tracking.

**Headline:** Automated drone-based animal tracking

## 1. Introduction

Technology has revolutionised our understanding of animal behaviour by enabling accurate collection of data across multiple scales (Patricelli, 2023). The global positioning system (GPS), in combination with radio telemetry sensors have shed light on mid- to large-scale spatio-temporal patterns of animal movement and social structure (Kays et al., 2015; He et al., 2023); bio-loggers collecting fine-scale data on animal posture and physiology, and have offered novel insights into cognition and social responses (Williams et al., 2020; Brandl et al., 2023); and passive monitoring devices such as acoustic recorders, camera traps, sonar imaging systems, and drones have revealed individual and collective behaviours of elusive species in a wide range of habitats (Burton et al., 2015; Nowacek et al., 2016; Munnelly et al. 2024). Recently, this development in electronic hardware has been coupled with the development of novel analytical and machine learning methods that can help recognize behavioural patterns and also inform the processes generating them (Valletta et al., 2017; Couzin & Heins, 2023; Christensen et al., 2024). One particularly powerful tool that is gaining more prominence is unmanned aerial vehicles (UAV), or drones. However, to date, drones have had limited application for tracking and georeferencing animals simultaneously, mainly due to challenges in analysing large video datasets.

Owing to their compact size, low cost, and high-resolution sensing capabilities, drones are finding increasing use in the study of free-ranging animals (e.g., Schad & Fischer 2022). Multi-rotor drones can be deployed from a stable platform in any terrain and provide detailed overhead perspectives on animals at different scales. Deep learning algorithms have been co-evolved to rapidly process drone imagery and provide accurate information on species identity (Bakana, Zhang & Twala 2024), individual counts (Eikelboom et al., 2019), body posture and behaviour (e.g., Koger et al. 2023; Dujon et al., 2021). Combined with photogrammetric methods, drones can also provide information on animal size and body condition (Dawson et al., 2017; Bierlich et al. 2024a), which can be used to infer their health (Ramos et al., 2022). Drones therefore have wide applicability in the study of health, behaviour and ecology across diverse vertebrate taxa, from fish (e.g., Butcher et al. 2021) to primates (e.g., Kays et al., 2019) to whales (e.g., Torres et al., 2020). Of their many applications, drones are particularly useful in conducting focal animal (or group) follows and record the interactions between animals and their environment or other conspecifics (e.g., Kays et al., 2019; Hartman et al., 2020; Brown et al., 2022) providing a perspective that is difficult or costly to achieve with conventional methods used in land- or boat-based surveys (Duporge et al., 2024), such as by fitting GPS tags to all group members (Grout et al., 2024; Papageorgiou, Nyaguthii & Farine 2024). However, analysing drone-based individual or group follow data is challenging as it often requires manually processing and annotating videos to describe behavioural patterns (Hartman et al., 2020).

Automated tracking of animals (Rathore et al., 2023) now also makes it possible to extract movement tracks—in pixel coordinates—of animals from drone video frames. These data can then be used to infer directional movements and behavioural states through Hidden Markov models, State-space models, and related techniques (Patterson et al., 2008; Patterson et al., 2017). However, positioning the animals in real space is more challenging. Current methods require the drone to remain stationary, and are constrained by requiring the animal movements to be within the bounds of the recorded video. Koger et al. (2023) recently solved this problem by developing a Structure-from-Motion-based approach (Jackson et al., 2020) to quantify landscape structure using overlapping drone images and then to translate individual pixel coordinates to obtain GPS tracks. While this approach provides highly accurate animal positions, it requires high computational power, a heterogeneous landscape, and ground control points (GCP) to geo-reference the landscape model. In addition, it requires the drone camera to be facing vertically downwards (i.e., in a nadir view).

The stringent requirements when using existing Structure-from-Motion solutions limit the applicability of tracking animals from drones across diverse fieldwork conditions. It remains challenging to track animals with drones in uniform habitats (e.g. open plains, at sea) and across different environmental conditions (e.g. different light conditions). Processing and analysing data is also time consuming. Further, methods cannot be applied in situations where the drone camera cannot face directly downwards. For example, at sea, the uniform surface precludes quantifying landscape structure, flying the drone directly overhead the animals may not be possible if propeller noise disturbs them (Raoult et al., 2020; Schad & Fischer 2022) and the high surfaces reflectivity and strong winds can make it impossible to film directly from overhead.

To overcome these limitations in drone-based animal tracking, we develop a tracking and geo-referencing pipeline to extract GPS-like tracks from drone videos of animals. Importantly, our algorithm relaxes the requirement of flying the drone directly overhead of the animals being tracked to accurately geo-referenced their position. We confirm the performance of our geo-referencing algorithm through multiple flights with varying drone and camera positions targeted at objects with a known georeferenced location (or Ground Control Point). We further validate the algorithm predictions by tracking a volunteer individual equipped with a high-resolution GPS unit and demonstrate the utility of our methods using drone videos of Lahille’s bottlenose dolphins *T. truncates gephyreus* and Indian Ocean humpback dolphins *Sousa plumbea* from independent studies in Brazil and India. These empirical exemplars capture varying environmental and flight conditions, thereby providing opportunities to discuss the advantages and shortcomings of our proposed workflow, as well as proposing general guidelines for drone use across different systems. Finally, we provide the necessary, and user-friendly, computational scripts (i) to obtain the geo-spatial position of any object in the drone video for cases that require manual tracking of animals, and (ii) to generate data through simple mouse clicks for re-training our algorithms for tracking animals in other systems.

## 2. Methods

Our analytical pipeline incorporates three components. The first is to detect objects of interest (e.g., an individual animal) within the video frame of the drone, for which we followed the MOTHE framework by Rathore et al. (2023). The second involves georeferencing the pixels in the video (i.e. the location of the detected objects within the frame) using the drone’s onboard GPS data and camera metadata, for which we developed a novel algorithm. The third involves linking objects across multiple frames to generate tracks, for which we used a Hungarian tracking algorithm. All analyses were performed in Python with post processing of GPS tracks in R (available at Supplementary Material S2).

### 2.1 Detecting objects

MOTHE (Rathore et al., 2023) uses a two-step R-CNN (Region-based Convolutional Neural Network) approach to first localise regions of interest (RoI) and then run a CNN model in these regions to detect the presence of objects. RoI are obtained by converting each frame into grayscale and then using a range of pixel intensity thresholds to identify ‘blobs’ using the OpenCV package (Bradski, 2024). These blobs can be filtered down based on their expected size and shape to increase computational efficiency. The approximate centre of each blob is called a ‘keypoint’ or anchor point, around a specified distance of which predictions are made by the detection model. To develop detection models to our empirical study cases (section 2.6), we trained a four-layered binary classifier CNN for Humpback dolphins that used animal and background images as training data; and we tracked Bottlenose dolphins manually.

The four convolution layers comprised of 64, 96, 128, and 256 filters respectively with a kernel size of 3×3 and MaxPooling layers of 2×2 between convolution layers. Each layer extracts increasingly abstract features from the image, ranging from simple edges and texture to complex shapes, which are used for image classification (Alzubaidi et al., 2021). The final output was flattened into a 1D vector and passed through two dense layers with a dropout rate of 30% to avoid overfitting. 60% of data was used to train the model while 40% was used to validate it. The model was trained for 25 epochs with a batch size of 64 and was evaluated using the accuracy, recall, and loss metrics (Alzubaidi et al., 2021). Training images were of the dimension 75×75×3 (width x height x RGB values) and were obtained for all datasets from the videos by manually clicking on animals and the background to extract their images in each video. We used data augmentation techniques to balance the number of animal vs background images for increased model performance (Huang et al., 2019), where new images were created by transforming (rotating, blurring, or jittering) dolphin training images to match the number of background images. We extracted dolphin images captured from different angles of the drone camera to allow their detection in any camera view. Models were trained to optimize performance based on categorical cross-entropy loss and using an Adam optimizer.

### 2.2 Georeferencing objects in drone videos

To geo-reference tracks we needed information on (i) drone flight parameters (height, orientation, and camera (gimbal) angle, (ii) camera lens parameters (focal length or field of view; FOV), (iii) video parameters (resolution, frame rate, frame number), and (iv) pixel position of each detection in the frame. All our videos were recorded using DJI drones that provide a log file for every flight. We used the DJI Flight Log Viewer (PhantomHelp, 2024) to convert the raw log file into a tabular, csv data file containing all flight information. The DJI website provides information on every model’s camera lens i.e., focal length and field of view (FOV), but since videos are recorded in a different aspect ratio compared to images, we estimated the horizontal and vertical camera FOVs for each of our study cases. We conducted an experiment where the drone was placed at a fixed distance (*d*) from a wall. The width (*w*) and height (*h*) of a part of the wall which was seen in the drone video were recorded. This was repeated 5 times and the drone’s horizontal and vertical FOVs were estimated, respectively, as:

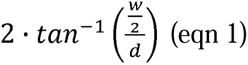

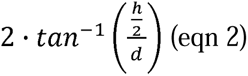

Frame rates were obtained from video metadata and frame number and pixel positions of detected objects in the frame were obtained in the previous step (section 2.1).

We synchronised all these data using timestamps that were available independently in each dataset. Timestamps for the video frames were obtained from the video’s SRT file or by adding frame time (frame number/frame rate) to the start time of the video. Timestamps for flight logs were obtained by adding flight start time (extracted from log file name) to flight time. Both the flight log time and the frame time were in the order of milliseconds, allowing a precise level of synchrony. However, in some cases flight parameters lagged by 1-4 seconds after synchronisation. To overcome this, we manually looked at 2-3 timestamps in the video where drone or camera movement began, and in the flight log where the same movements were recorded. We then estimated lag time and synchronised the two files with the defined offset.

To geo-reference any object, we need information on how far (*D*), and at what angle (or azimuth) (Φ), the object is from the drone. The distance of an object at the centre of a frame can be estimated if the height (*h*) of the drone, and the angle of its camera with respect to the ground (θ) is known, as D = h · tan(0). However, for an object away from the centre, these parameters need to be explicitly estimated. In other words, how much should the drone’s camera angle (θ) and drone’s orientation (Φ) change to bring the object to the frame’s centre? (Fig. 1).

**Fig. 1:**
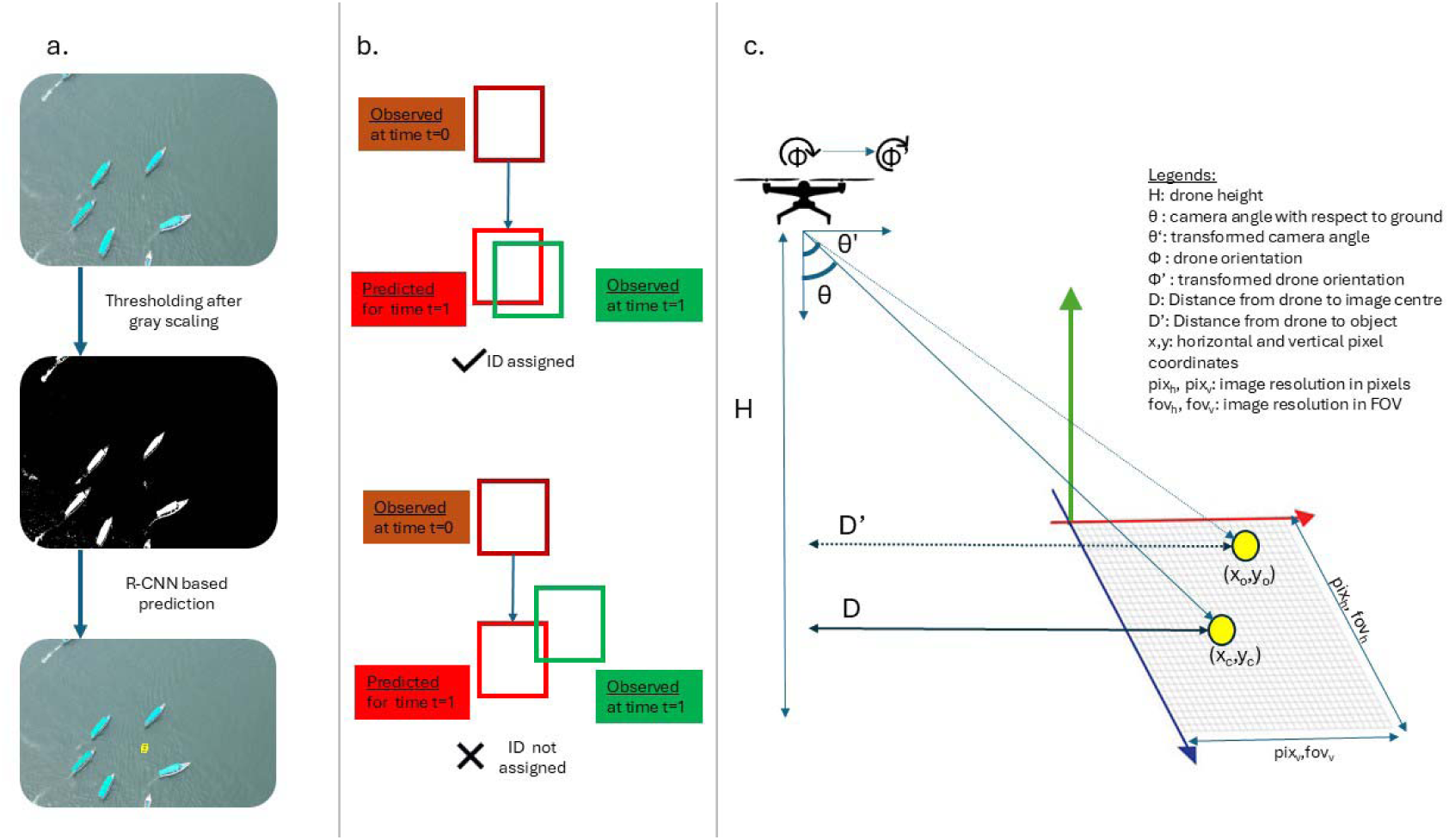
A schematic of the process involved in automated tracking of animals. An R-CNN network is used to locate animals in a frame (a) followed by a Kalman filter based approach to link detections across frames (b). Animal tracks are then georeferenced based on flight parameters and their pixel location on the frame (c).

To answer this, we first scaled the pixel position of the object in terms of the camera’s horizontal and vertical FOV as:

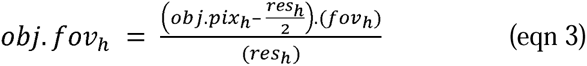

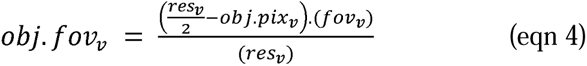

where *obj.fov* is the angular position of the object (detection), *obj.pix* is the pixel position of the object, *res* is the image resolution, and *fov* is the camera FOV when recording; *h* and *v* represent horizontal and vertical components respectively. The parameters were then estimated as:

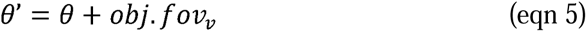

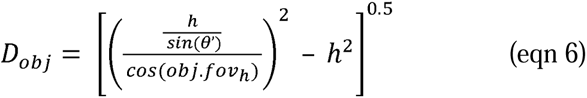

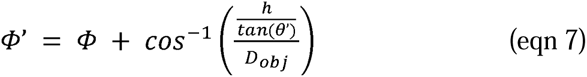

where θ*’* is the adjusted camera angle*, D_obj_* is the distance of the object from the drone, and Φ*’* is the adjusted drone orientation. The object’s latitude and longitude can be estimated as function of these parameters and the drone’s geo-spatial position, as available from the flight log.

### 2.3 Linking objects to generate tracks

Each detection in a frame was provided with a unique identification code (ID), and its position in the next frame was predicted based on a Kalman filter; these positions were linked using a Hungarian algorithm (Hamuda et al., 2018; Rathore et al., 2023). The Kalmann filter reads the pixel position of an object and estimates its speed, acceleration, and direction to predict its location in the following time frame. Since each object is defined by a bounding box, the Hungarian algorithm estimates and minimizes a cost matrix for assigning IDs to new detections based on their overlap with the predicted locations of all previous detection. By combining these, we were able to detect and track individuals across the frames of each video.

It is challenging to detect all animals in a single frame, especially in marine environments where animals can occur in tight groups (Saqib et al., 2018). Therefore, to improve our accuracy, we used the total number of unique tracks in a small-time window (4 sec) instead of a single frame to estimate animal count (Brown et al., 2023).

### 2.5 Ground-truthing the algorithm

We tested our geo-referencing model by conducting eight independent flights where 1-2 points with known geo-coordinates were viewed from different flight heights (from 15 to 120 m), drone angles (from -90° to 0°) and horizontal distances (from 0 to 300 m). Error was calculated as the distance between estimated coordinates and true coordinates. We also conducted two flights to estimate the size of a known linear object by estimating the geo-coordinates of its two ends in the same format. We used GLMs to understand how drone height (*h*), object horizontal position in frame (*obj.fov_h_*), and adjusted camera angle (θ*’*; estimated from object vertical position in frame; section 2.2) impacted error rates for geo-referencing and size estimation.

### 2.6 Empirical applications

To first demonstrate the ability to reconstruct tracks from moving animals, we followed a volunteer individual from our team carrying a high-resolution GPS unit. We set our drone to fly at an altitude of <50 meters from the ground and estimated their track using our methods (section 2.2). We plotted the tracks from both the GPS unit and estimated from the drone to visualise error in track generation.

To demonstrate the real-world utility of our methods, we used drone videos from 3 empirical case studies that represent varying environmental and flight conditions (Table 1). The first case study is of Lahille’s bottlenose dolphins (*T. truncatus gephyreus*) and their interactions with net-casting fishers in the shallow waters of Laguna, Brazil (Cantor, Farine & Daura-Jorge, 2023; Fig. 3a). In an exemplar drone video, a group of 3-6 dolphins forages on mullet schools near the edge of a steep canal where fishers wait for the dolphins’ cue that they interpret as the right time to cast their nets over the mullet (Supplementary Material S1). The dolphin-fisher interactions were recorded from a nearly stationary drone at a fixed elevation with minimal drone or camera movement. The other two case studies are of Indian Ocean humpback dolphins (*S. plumbea*) recorded in the coastal waters of Goa, India. In the second case, the drone was first flown at a fixed altitude of 98 m to follow a large group of dolphins consisting of 6 to 8 subgroups that vary in size—as individuals split, joined, and interacted with each other—for a duration of 1 minute 45 seconds (Fig. 3d). The drone flight trajectory, while fixed in height, intermittently moves along with the dolphins (Supplementary Material S1). In the third case, a group of slow-moving dolphins in 2-3 sub-groups with 4 to 5 satellite individuals around them were followed for 22 seconds; a single dolphin from these satellites approached and joined the central group towards the end of the bout (Fig. 3g). The drone was flown at a similar altitude as the first case (98 m) but with higher levels of drone and camera movements (Supplementary Material S1).

**Table 1:** Summary of the three drone videos and drone flight parameters used in this study.

| Species | Location | #groups | #individuals | Drone | Tracking duration (s) | Drone height (m) | Drone camera angle | Camera horizontal FOV (°) | Camera vertical FOV (°) |
| --- | --- | --- | --- | --- | --- | --- | --- | --- | --- |
| Bottlenose dolphin | Laguna, Brazil | - | 6-8 | DJI Phantom 4 pro V2 | 981 | 70-80 | -80 to -90 | 70 | 40 |
| Humpback dolphin | Goa, India | 6-8 | 70-80 | DJI Mini 3 pro | 105 | 90-100 | -60 to -70 | 70 | 40 |
| Humpback dolphin | Goa, India | 2-3 | 20-22 | DJI Mini 3 pro | 22 | 60-70 | -40 to -70 | 70 | 40 |

For humpback dolphins (case studies 2 and 3), we first built a detection model (which had high accuracy and recall, Table 2). We then georeferenced and tracked individuals to estimate the number of animals, their movement speed, direction, and body size. These case studies illustrate one major utility of our method, which is to help overcome the challenge of estimating numbers when individuals are asynchronous and/or highly clustered in their behaviour. In the videos, several dolphin sub-groups were visible from a high altitude simultaneously and individuals were spaced closely which could result in incomplete detection (Saqib et al., 2023) and loss or switching of identity during tracking (Rodriguez et al., 2017). To accurately identify groups and their tracks, we performed a time series spatial-clustering analysis on extracted tracks using the *dbscan* package (Hahsler, Piekenbrock & Doran, 2019) in R. A sub-group of dolphins then consisted of one or more individuals within 10 meters of each other (Syme, Kiszka & Parra, 2022). A change in group size (defined as several sub-groups within the area) between sub-groups was allowed. Single individuals were also identified similarly. IDs that were assigned to individuals but not detected for at least the next 60 frames (2 seconds) were likely false positives and were discarded. Of the remaining IDs, each detection of an individual ID, was assigned a sub-group ID and its track could therefore be mapped as ‘a sub-group’ in case of loss of the individual ID. As no individual can be a part of two sub-groups at the same time, we used this information to estimate (i) change in the number of dolphin detections throughout the video, and (ii) cumulative number of dolphins detected. We also manually counted the ‘true’ number of dolphins present in the videos and compared it with these estimates. To avoid error in estimation due to drone or camera movement, we measured speed of movement only when the drone was mostly stationary (<5 km/h).

**Table 2:**
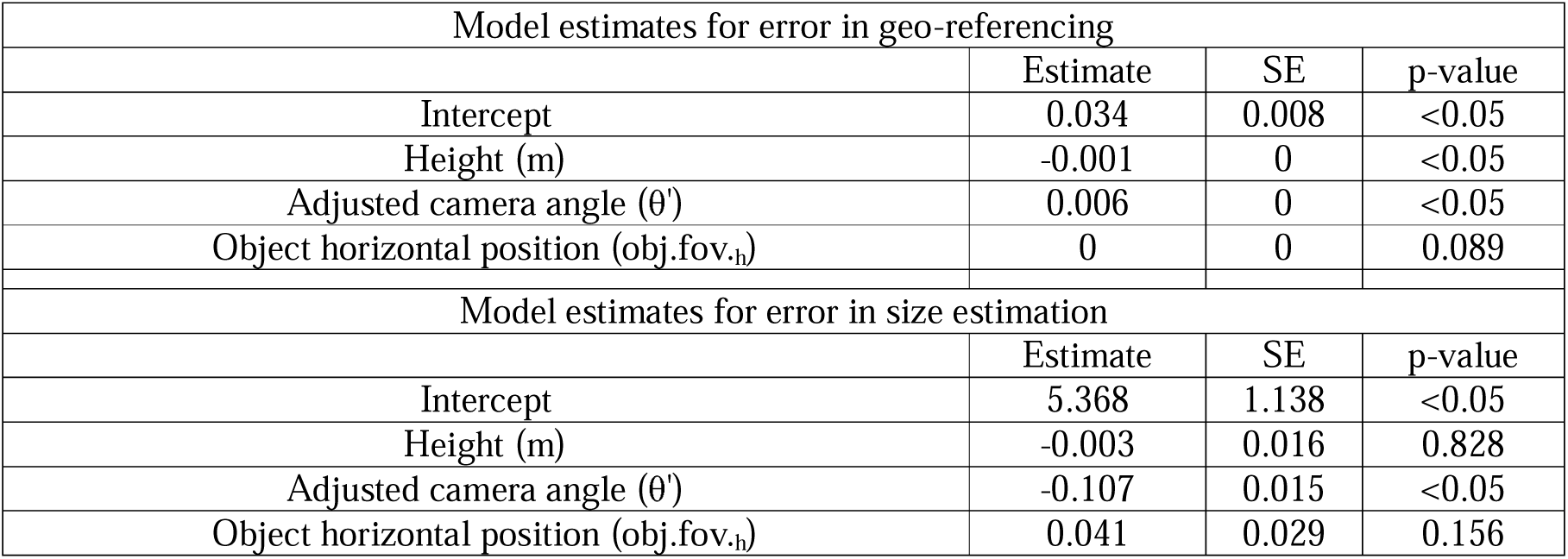
Model summaries for error in geo-reference and size estimation. Geo-referencing error (m) was modelled using a GLM with a gamma error distribution and inverse link while size estimation (%) was modelled using a linear model.

Since bottlenose dolphins surface rarely when interacting with cast net fishers (case study 1) and are only partially visible in the murky water in each surfacing (Cantor et al. 2023), we tracked individuals manually for a complete flight duration of ∼17 minutes and estimated their movement (distance from fishers over time using our geo-referencing model.

## 3. Results

### 3.1. Validating geo-reference models using ground-truthed data

We tested our geo-referencing model with several independent flights. The median error in estimating geo-coordinates was 2.8 (0.7 – 23.2; 95% CI) m (Fig. 2). Our data (n=762) suggested that error increased by 0.1 cm and 0.6 cm with unit increase in drone height (*h*) and adjusted camera angle (*θ’*) respectively but was not related to the horizonal position of the object in the frame (Table 2). Similarly, the median error in estimating object size was 6.7% (0.8-17.7%) which decreased by 0.1% with an increase in adjusted camera angle (*θ’*) by 1 degree. However, error did not change with drone height or orientation (Table 2).

**Fig. 2:**
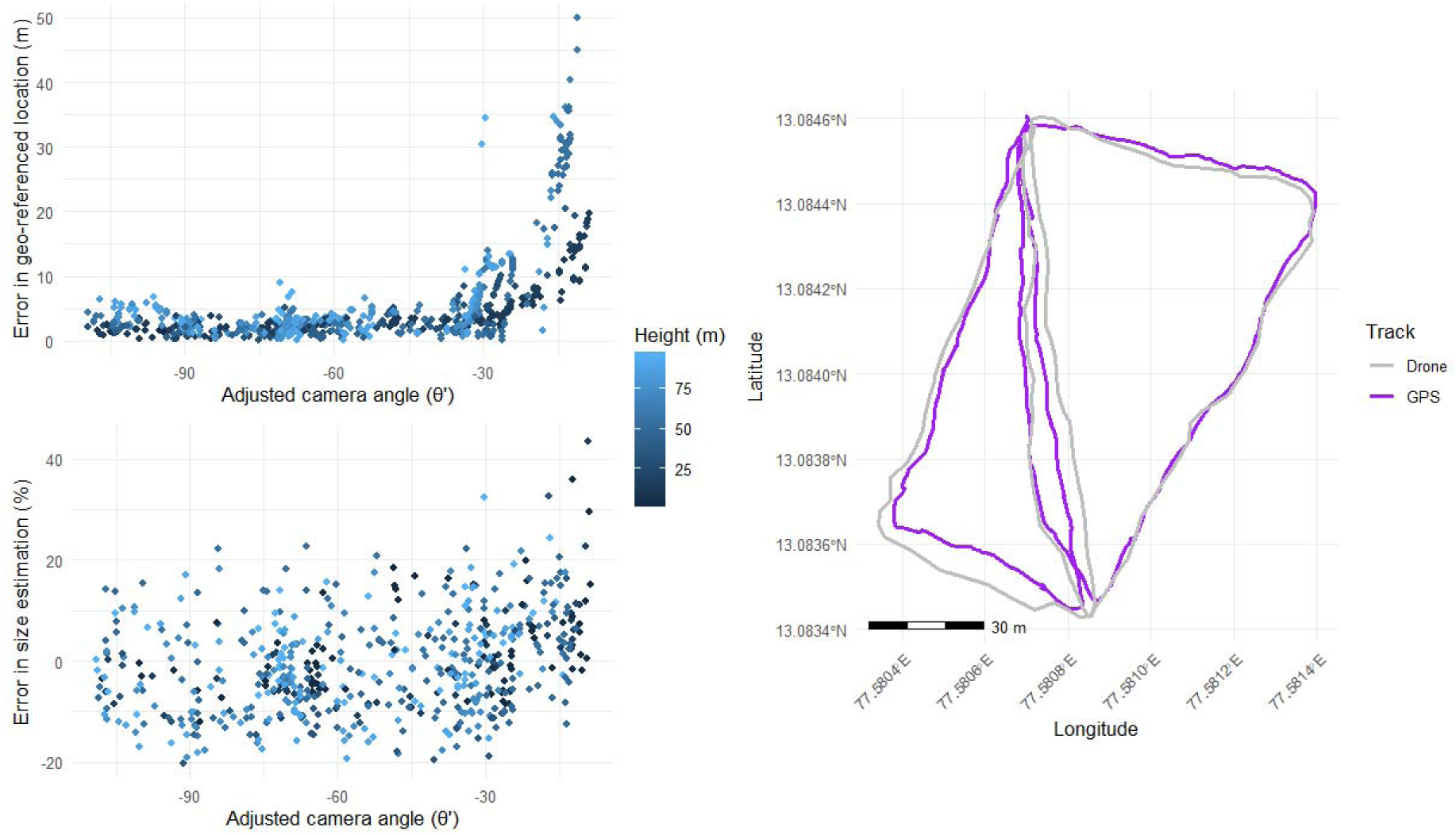
Impact of drone flight height and camera angle on estimation of geo-coordinates (top) and size (bottom). An angle of 0° refers to the camera pointing at the horizon while -90° refers to the camera pointing straight downwards (nadir view). Adjusted camera angle refers the drone camera angle when the object is at the frame centre. The right panel represents volunteer tracks recorded in a GPS and estimated from drone footage.

Geo-reference errors were minimum (mean: 1.5, 95% CI: 0.5 – 2.8 m) when the camera was positioned facing vertically downwards (nadir view) and the drone was flown at <50 m. This error did not increase significantly as long as the camera angle was kept below -40° (see Supplementary Fig. A1 in S1).

### 3.2. Reconstructing animal tracks

Our algorithm was capable of reconstructing the GPS track of a moving volunteer with high accuracy. The estimated track from the drone overlapped closely with tracks from the GPS unit they carried (Figure 2), producing a mean error of 2.38 (0.47 – 5.63; 95% CI) m, which is only slightly elevated from the error from fixed points.

#### Case study 1: Tracking individual bottlenose dolphins foraging with humans

We estimated 12 dolphin tracks based on relative surfacing positions and timings of individual Lahille’s bottlenose dolphins. Only 6 out of these had more than two surfacing events and were plotted on a map. Dolphins regularly moved in and out of the video frame and so it was impossible to estimate the actual number of dolphins in the area based on drone images alone without implementing an algorithm to individually recognise dolphins.

However, field observations and photo-identification based on dorsal fin marks carried out from shore (details in Cantor et al. 2023) suggest there were 5, maximum 6, individuals foraging with the fishers during the whole 17-min video. Fisher positions were accurately geo-referenced, and so were dolphin tracks obtained from object detections where we could establish individual identity with certainty based on their dive orientation and dive times (Fig. 3b). Dolphins generally foraged >30 m away from shore but approached fishers to as close as 12 m midway through the recording (Fig. 3c). Fishers were seen casting their nets following the foraging cues of approaching dolphins (Supplementary Material S1).

**Fig. 3:**
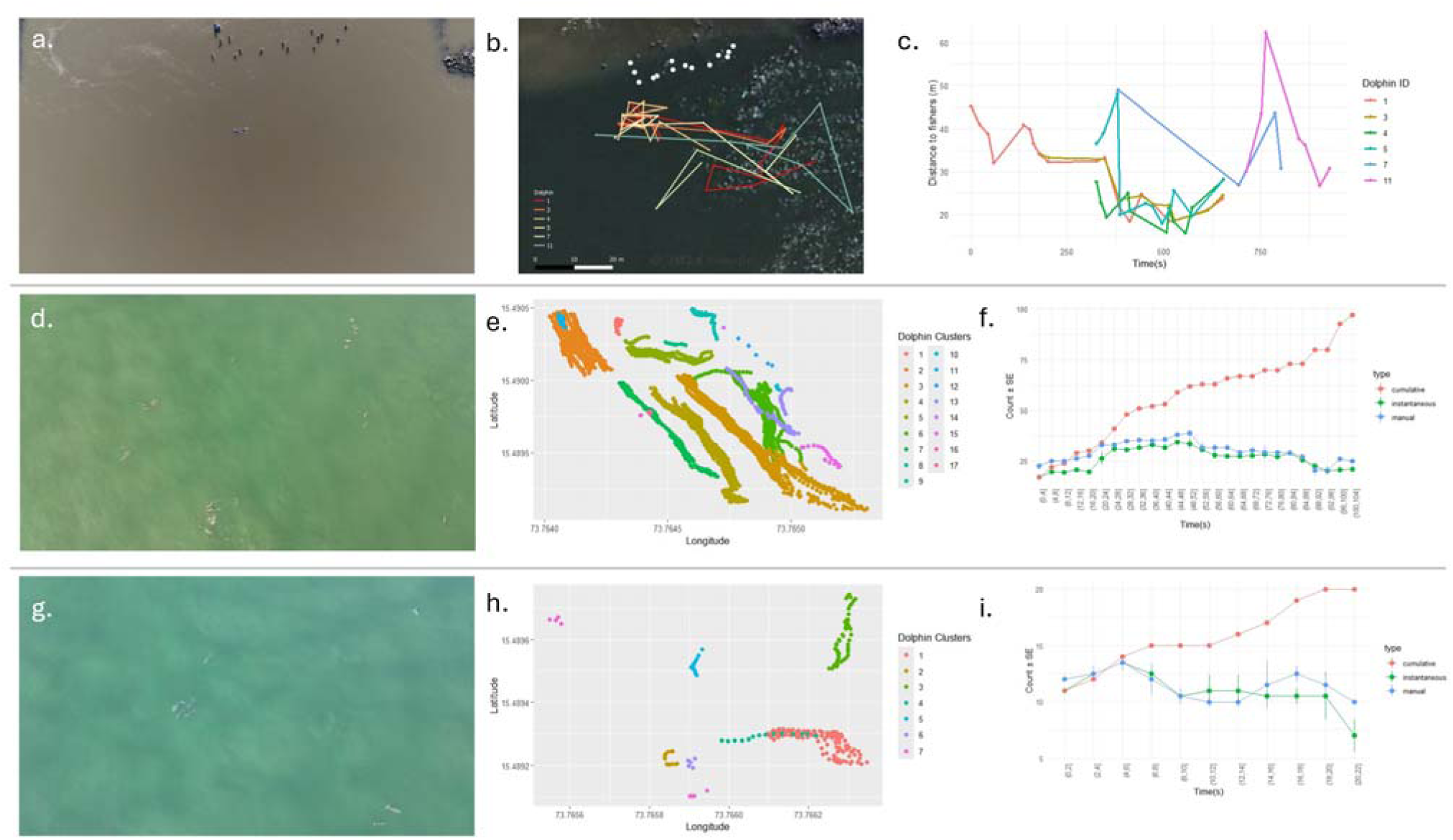
Drone video snapshots (a, d, g) estimated tracks (b, e, h), and animal counts (f, i) for the two species, Lahille’s bottlenose dolphins and Indian Ocean humpback dolphins (cases 1, 2, and 3). Distance of bottlenose dolphins from fishers overtime is show in panel ‘c’.

#### Case study 2: Tracking large humpback dolphins groups nearshore

Indian Ocean humpback dolphin detections throughout the duration of the video (1 minute and 45 seconds) ranged between 17-36 individuals (Fig. 3f) spread across in 17 sub-groups (Fig. 3e). The cumulative number of dolphins were estimated at 97 individuals but since detections were underestimated by an average of 11.4% (± 8.9 SD), the true number of individuals present in the group was likely higher (Supplementary Material S1). The dense clustering of individuals combined with drone movement led to a high number of false detections (detections lasting for <2 seconds in total), which were removed during subsequent analysis. Individual IDs were consistent during tracking when the drone and/or dolphin movement was minimal but were lost at several instances. Therefore, we were not able to track individuals consistently over time. However, we identified and successfully tracked 17 sub-groups with a mean size of 6.1 (± 5.1 SD) individuals. Mean dolphin length was estimated at 164 (± 35 SD) cm and did not vary significantly across sub-groups. The mean sub-group speed was 0.93 (± 0.48 SD) m/s (Supplementary Fig. A2 in S1).

#### Case study 3: Tracking small humpback dolphins groups with drone and camera movements

The second video of Indian Ocean humpback dolphins was shorter (duration 22s; Supplementary Material S1) but involved a greater movement of the drone and camera (Table 1). We detected two clusters and five single individuals with a cumulative dolphin count of 20 individuals (Fig. 3i). Detection bias was relatively low (4.3 ± 11.7 %) because of lower flight height, and more importantly, neither drone nor camera movement impacted predicted tracks of the individuals/groups (Fig. 3h). Similarly, individual IDs were also more consistent throughout the video with a lower rate of incorrect detections.

## 4. Discussion

We present a novel geo-referencing algorithm combined with deep learning methods for object detection to obtain highly accurate, GPS-like tracks of individual and groups of animals from drone video data. We demonstrate that animal tracks can be obtained under challenging field conditions (e.g., over the uniform, glistening sea surface) and across any combination of drone flight heights, camera orientation and angles, while ensuring track accuracy despite drone or camera movements. Our proposed methods and workflow resolve the need to fly drones directly over animal groups with the camera in nadir, and allows drone pilots to adjust their flights according to a wide variation in environmental conditions and animal behaviours.

### 4.1. Advancing drone-based animal behaviour studies

Drones are now widely used in surveying biodiversity (Lyons et al., 2019; Corcoran et al., 2021) and estimating animal abundances (Eikelboom et al., 2019) as their aerial perspective allow the remote observation of animals with a high level of detail (Hartman et al., 2020). Therefore, drones can provide the basis for understanding animal behaviour and its social or environmental drivers with minimal, if any, disturbance to groups when flying at appropriately high altitudes (Schad & Fischer 2022). Drone model, operations, and usage must be defined by the study’s specific objectives, the environment, the focal species and the local flight regulations. For example, species differ in their sensitivity to responses to drones (Schad & Fischer 2022) and so the drone model must be appropriately chosen or modified to minimise noise from rotors (Raoult et al., 2020). Similarly, groups that are spread across areas larger than what can be covered by the drone camera’s field of view may require multiple drones to be flown in parallel. Data from all these drones can be processed independently using our framework to obtain tracks of all individuals. Alternatively, the drone can be positioned with an oblique camera angle that captures all animal groups, but making the angle too steep (>-40°) can come at the cost of increased geo-referencing errors.

Unlike a Structure-from-Motion approach (Jackson et al., 2020), our proposed framework does not require information of habitat features and is therefore useful for a wider range of applications. This includes tracking animals in homogeneous environments (like the sea or on snow) or at night when combined with thermal cameras. Accurate georeferencing also opens the door to collecting additional pieces of information, for example the size of horizontal features such as boats near dolphins can be obtained by capturing two ends of the object. A major advantage of georeferencing animals is the ability to collect continuous data on individual animals. Drone batteries typically constrain fly time to less than 40 minutes, thereby limiting the length of aerial observations that can be recorded. If longer behavioural observations are needed, a second drone can be launched right before the first drone is set to retreat (see Koger et al. 2023). Since our tracking algorithm is independent of the drone position, uninterrupted information can then be obtained by stitching animals tracks and identities from the two drone videos together.

### 4.2. Caveats of current methods

#### 4.2.1. Consistent identity assignment during tracking

Our primary aim was to develop a georeferencing algorithm for obtaining information such as the location and size of objects from drone images. We also explored the potential to combine these data across many images to produce GPS-like tracks of animals, with a specific interest in animal groups. Like with previous tracking algorithms (Rodriguez et al., 2018; Sridhar, Roche & Gingins 2019), our models had limited success in consistently tracking individuals when animals were tightly grouped or remained submerged for extended periods. Individual humpback dolphins could be tracked when the drone was stationary, but their identities were more likely to be lost or misassigned when there was drone movement or when dolphins interacted too closely with one-another. In such cases, it is difficult to track individuals manually as well, and so changing tracking-model parameters such as the bounding box size or Intersection over Union (IOU) thresholds may not be very useful. Thus, if it is essential to preserve individual identities, then we recommend minimising the amount of drone movement. Flying at lower altitudes can also help, but this comes with a trade-off of spatial coverage. In case where both are required, one drone could be flown at low height to consistently track individuals and another flown at a higher height to record other groups/structures. Nevertheless, our application may still be prone to errors in identity assignment and tracking, especially if flight parameters are not fully synchronised (section 2.2) and we recommend crude verification of animal tracks before they are used for further analysis.

Advances in machine learning algorithms are rapidly improving our ability to collect data from animals. For example, along with animal tracks, information on animal posture can also be obtained by mapping the relative positions of different body parts of each individual (Pereira et al., 2022). An important application of our work is in the study inter-group interactions, because each group can be described and tracked as a geographical cluster with known number of individuals. In cases where multiple species need to be tracked simultaneously, our R-CNN can easily be modified into a multi-class detector with minor changes to the model building code and downstream analytical pipeline. Finally, our method can only track animals on the ground surface. In cases where animals are moving across rugged terrains or changing elevation our geo-referencing algorithm will need to be modified to incorporate habitat structure through digital elevation maps.

#### 4.2.2. Training data and computational efficiency

An important requirement for building object detection models is good quality training data. The volume of training data required can vary depending on environmental complexity, animal size and other factors (see Rathore et al., 2023). We provide a simple computational script to obtain training (animal vs background) data of desired size from the videos themselves (Supplementary Material S2). For common animal species, off-the-shelf detection models can also be downloaded from other sources (e.g., Bakana, Zhang & Twala 2024), or in simple cases, detections can be marked manually (section 2.6, case study 2). When automating detections, there are likely to be false positive and missing detections across different frames (Saqib et al., 2018), which can make it difficult to estimate animal counts. We overcome this issue by estimating the number of consistent tracks over a small time period rather than estimating animal counts for each frame.

An important issue with automated tools is computational power requirements. Our framework provides fast and accurate detections (processing ∼ 1 frame/sec), with the georeferencing part of the algorithm accounting for a small fraction of this time. However, the pipeline may become slower if a high number of animals are to be tracked. For such cases, our Region-based CNN can be replaced with a faster GPU-based YOLO model (Terven, Córdova-Esparza & Romero-González 2023) with minimal changes to downstream analysis. However, it may be inefficient if object sizes are too small and would require more specialised computational hardware.

#### 4.2.3. Varying drone quality and manufacturing

Our three case studies used videos recorded from readily available commercial grade DJI drones. The least expensive one was the DJI Mini 3 Pro that costs under USD 1,000 and weighs 249 g. Commercially available technology even in relatively cheap equipment is highly accurate (Hussein et al., 2022) but may sometimes be prone to errors. Altitude sensors, for example, rely on atmospheric pressure measurements to estimate drone height, and can provide imprecise estimates. One solution to this is to supplement data with laser-based altimeters that provide highly accurate measurements (Bierlich et al. 2024b). Similarly, drones can be supplemented with real-time kinematic (RTK) technology, which enhances GPS precision by using data from a nearby base station in addition to satellite signals, enabling centimetre-level accuracy. Since we used off-the-shelf drones for our work, some error in object position estimation may relate to inaccuracies in the drone’s sensing abilities. Therefore, better equipped drones are likely to provide a more accurate positional accuracy and size estimate for objects.

We also observed that flight parameters recorded in the flight log lagged by up to 4 seconds in some cases, which caused errors in geo-referencing because of erroneous synchronisation. While such errors may not occur in higher grade drones, we recommend checking synchrony for each flight at least once using our method.

## 5. Conclusion

Extracting individual animal tracks from drone videos can be used to systematically study fine scale animal behavioural responses to the environment, conspecifics, and other species. Our novel approach provides the flexibility of collecting such data independent of specific drone flight requirements (i.e., flying over animal groups). Animal tracks can be extracted quickly with high accuracy and without the need for high-end hardware. In addition, the modular structure of our framework allows easy modification of the different steps (e.g., detection algorithm) to adapt to specific cases, when needed. Our application will be useful in many cases ranging from understanding animal movement and behaviour to estimating body sizes and population age structure.

## Acknowledgements

This manuscript is part of I.S.’s PhD thesis at CES, IISc which was supported by the Prime Minister’s Research Fellowship. We thank H. Patil, P. Mitra, and all our project collaborators from Goa for their support during field work. We acknowledge support from the Goa Forest Department and the Flag Officer Commanding, Indian Navy at Goa in providing necessary permits to conduct this work. We thank the support and logistics from our colleagues when conducting fieldwork in Brazil, especially those directly involved in the drone sampling (F.G. Daura-Jorge, A.M.S. Machado, P.V. Castilho). I.S. received support from the Rufford Small Grant program (41484-1) for conducting field work in Goa. M.C. received support from the National Geographic Society (NGS-101549R-23) and the Marine Mammal Research Program Fund at Oregon State University.

## Data availability statement

All data and codes relevant to this study will be available in a public GitHub repository upon editorial decision.

