## Supplementary Material S1 for "Integrated deep learning and geo-referencing for drone-based animal tracking with flexible camera angles"

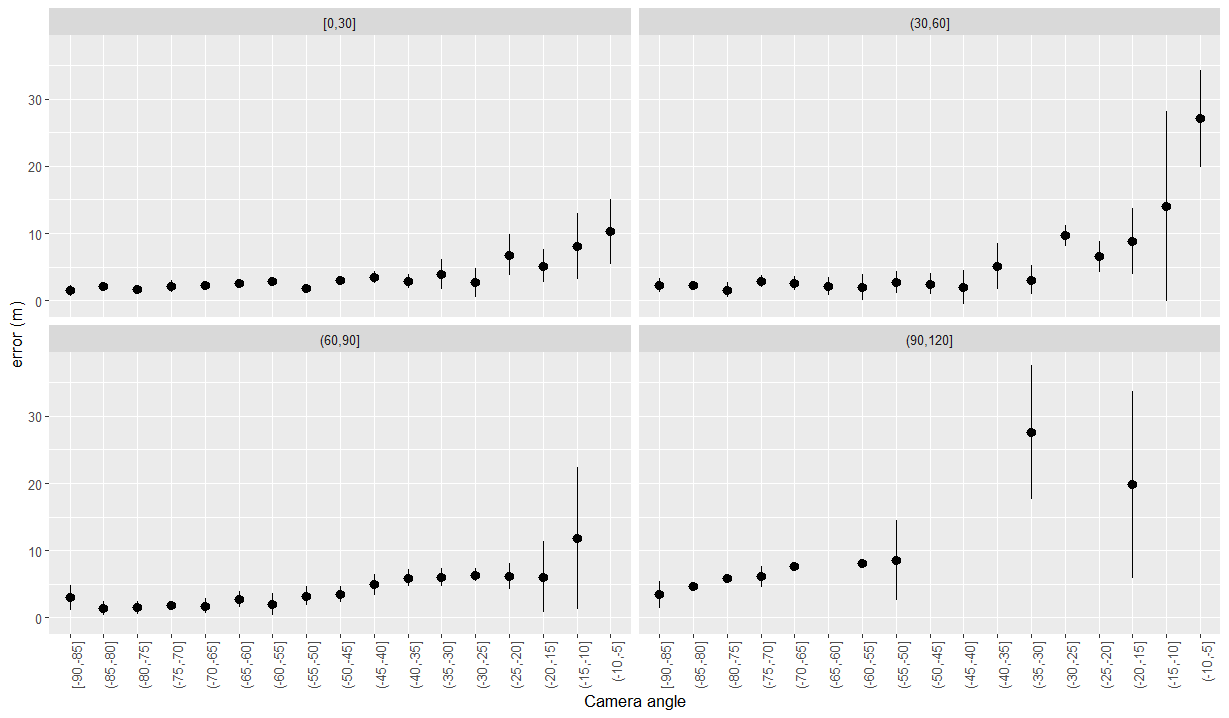

Figure A1. Error in geo-referencing vs camera angle for different drone heights

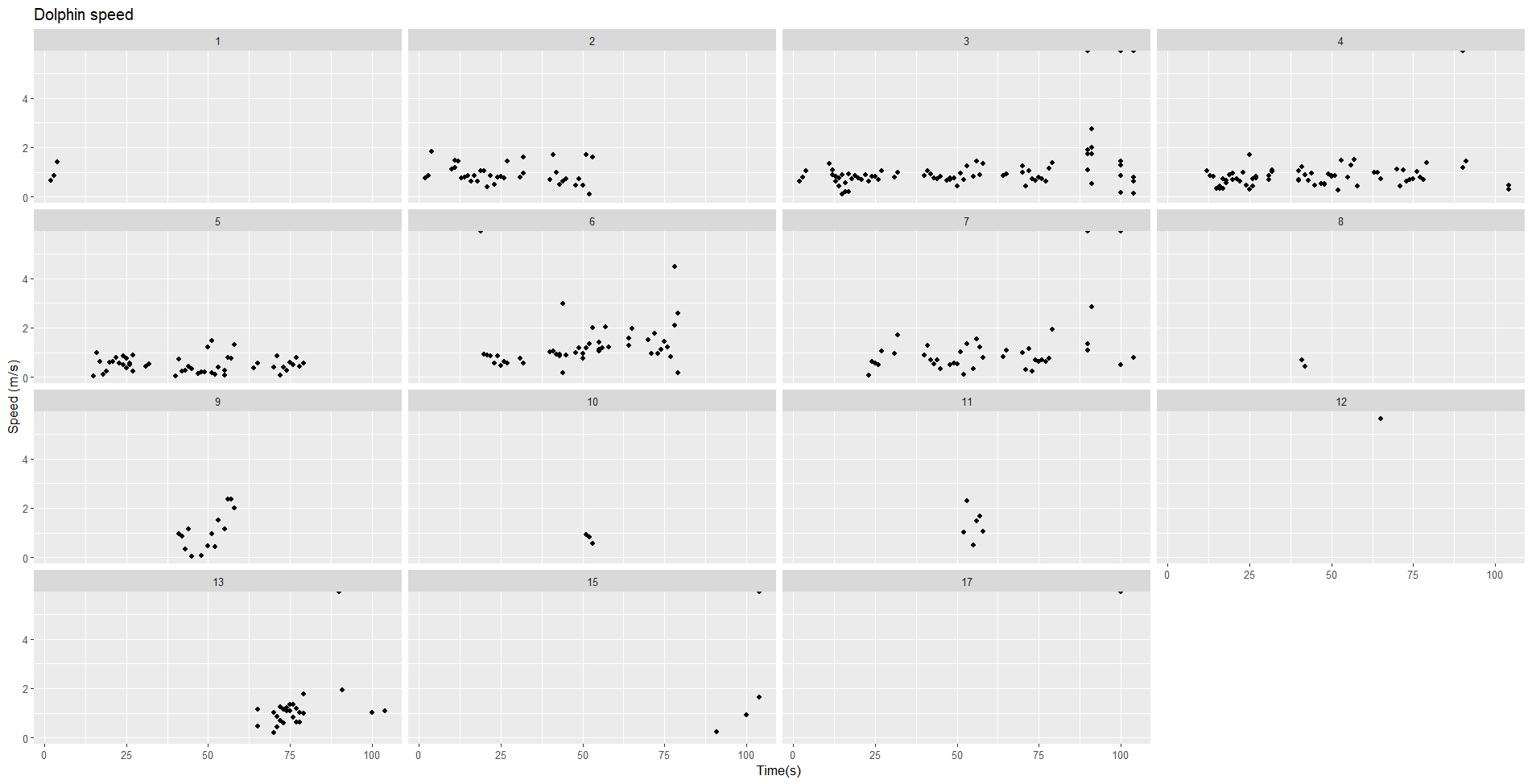

Figure A2. Estimated dolphin group speed vs time for different groups

The videos used in the manuscript can be found in this folder:
http://e.pc.cd/jN2y6alK
